# IsoAtlas: Visual interpretation of known and novel transcript isoforms using population-scale long-read evidence

**DOI:** 10.64898/2026.08.12.744438

**Authors:** Xinchang Zheng, Fritz J. Sedlazeck

## Abstract

Long-read RNA sequencing has revealed extensive transcript diversity, but newly observed isoforms remain difficult to interpret beyond their classification as known or novel. Here, we present IsoAtlas, an interactive multispecies database for visual exploration and population-scale interpretation of transcript isoforms using 1, 035 human and 414 mouse uniformly processed long-read RNA-sequencing samples. Users can search annotated genes and transcripts or submit novel transcript models in GTF format, visualize their structures, and examine sample-level support, prevalence, expression, tissue and disease context, and sequencing-platform evidence. IsoAtlas integrates structurally equivalent transcripts across GENCODE, RefSeq and CHESS, consolidating evidence that would otherwise be distributed across annotation-specific identifiers. It further links corresponding human and mouse transcript models, enabling users to assess cross-species conservation and enabling users to assess cross-species conservation and inform the suitability of mouse models for isoform-specific studies. IsoAtlas can also evaluate arbitrary user-supplied transcript structures directly against accumulated long-read evidence. IsoAtlas therefore complements established reference annotations with an extensible evidence layer that connects transcript structure to population prevalence, biological context and cross-species support. IsoAtlas is freely available at https://www.isoatlas.org/.

**Graphic abstract:** 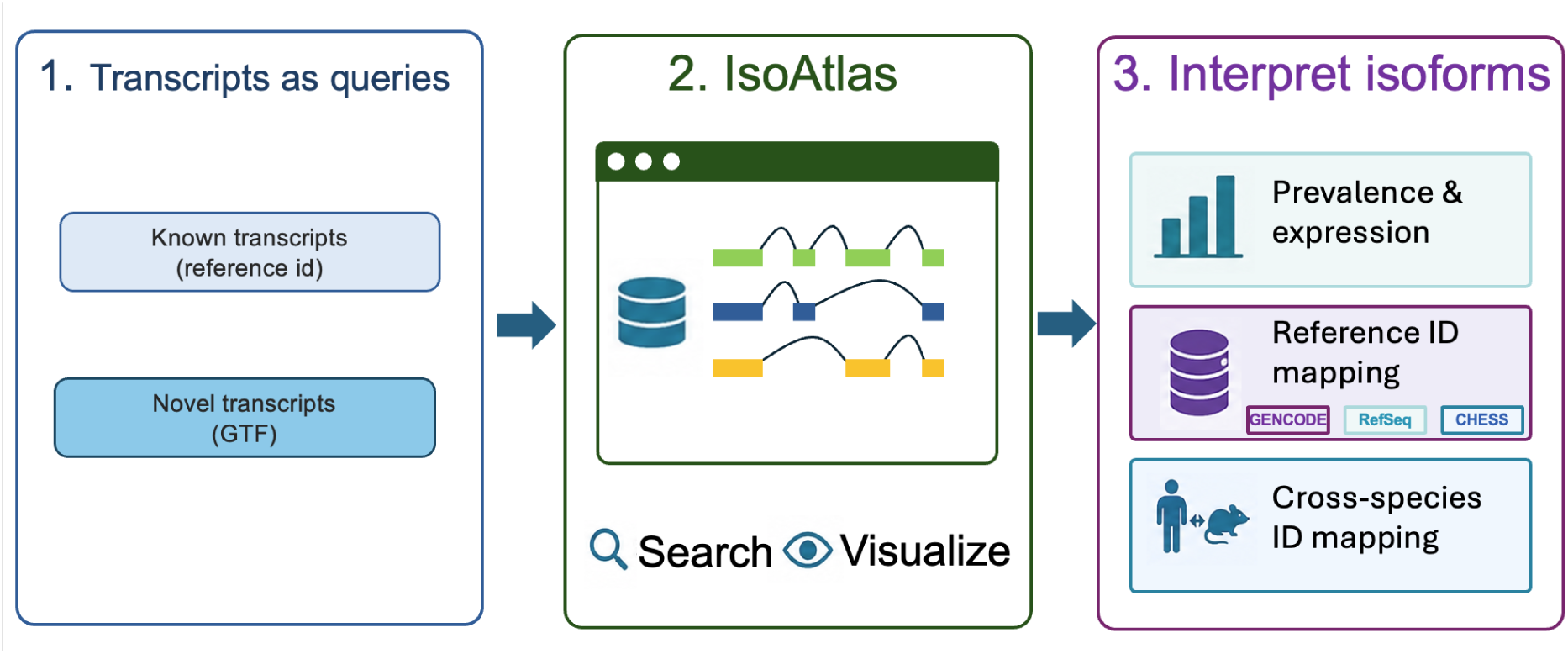

## Introduction

Long-read RNA sequencing enables direct observation of full-length transcript structures, including complete combinations of exons, splice junctions, and transcript ends within individual molecules(1, 2). This capability has substantially expanded our understanding of isoform diversity beyond that represented in static reference annotations. At the same time, long-read sequencing has exposed a fundamental limitation in the current interpretation of transcript novelty. Most long-read studies report large numbers of “novel” isoforms, yet novelty is defined relative to the specific reference catalog and version used for analysis(3). Consequently, an identical transcript structure may be classified as novel in one study but annotated in another. As reference catalogs continue to expand, transcripts previously designated as novel may subsequently be incorporated without any change in their underlying biology. Transcript novelty is therefore not an intrinsic or stable biological property, but a classification that depends on the analytical context.

Major reference resources, including GENCODE(4), RefSeq (5), MANE(6), and CHESS(7), remain essential foundations for transcriptomic analysis. However, they differ in transcript coverage, annotation strategy, evidence requirements, and intended use. Long-read studies provide complementary evidence, but their transcript models are generally distributed across individual publications and processed using different reference annotations and analytical pipelines. In addition, human and mouse transcript resources are typically examined separately, making it difficult to determine whether an isoform is conserved across species, represents a lineage-specific structure, or is supported in a model organism relevant to human disease.

Several databases integrate evidence related to alternative splicing and transcript diversity, including IntroVerse(8), RJunBase (9), TCGASpliceSeq(10), recount3(11), and AScancer(12). These resources are derived predominantly from short-read sequencing, which can quantify splice junctions and splicing events across samples but cannot directly resolve complete transcript structures within individual RNA molecules; accordingly, sample-level prevalence in these resources is reported at the junction or event level, and none supports querying by reference annotation transcript identifiers. Novel transcripts are often underrepresented in databases built exclusively from short-read datasets. FLIBase additionally incorporates long-read RNA sequencing data from 351 samples, enabling direct representation of full-length transcript structures, with isoform-level expression estimated by mapping short reads onto its long-read-derived models(13). However, its novel transcripts are indexed by internal catalog identifiers rather than reference annotation IDs, and per-transcript detection frequencies across samples are not explicitly reported. More fundamentally, all of these resources are organized around a fixed catalog of pre-computed features: none allows users to submit an arbitrary, user-defined full-length transcript structure and retrieve its population-scale sample prevalence, and none provides cross-species transcript correspondence between human and mouse. A direct feature comparison is available in **Table 1**.

**Table 1:**
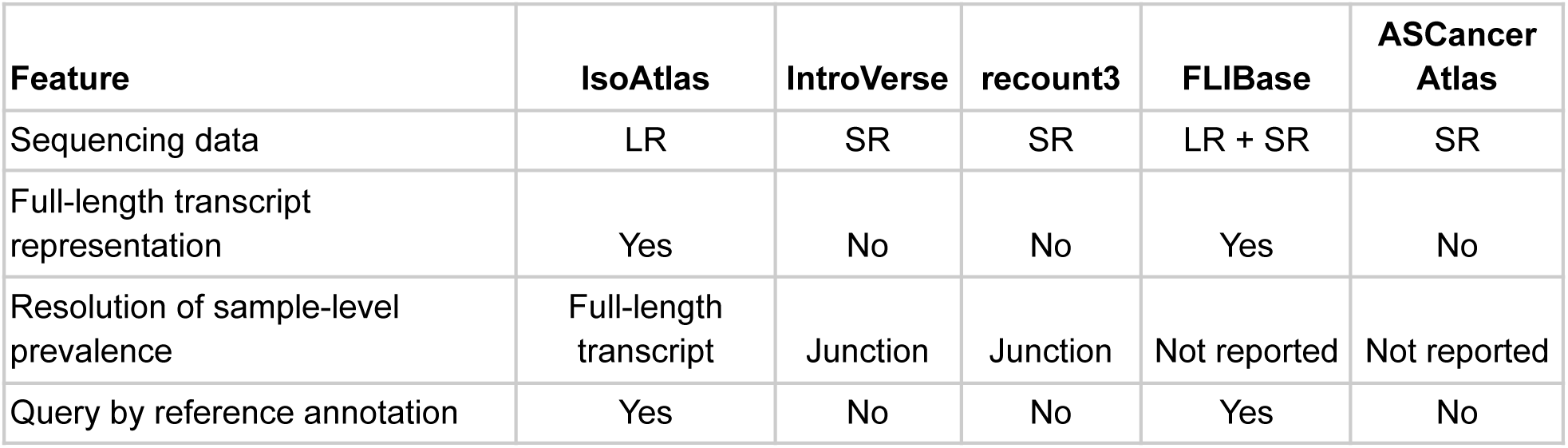
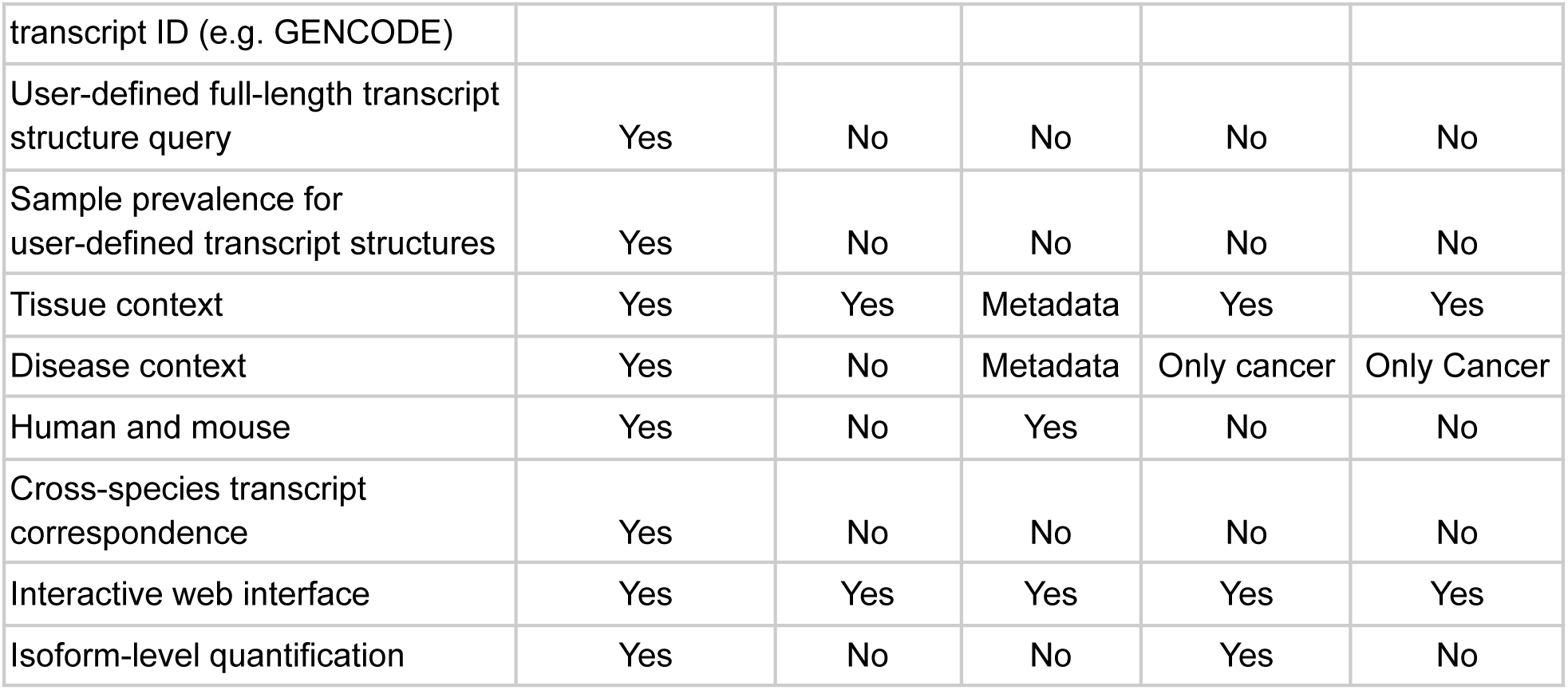
Feature comparison of IsoAtlas and published databases.

To enable direct exploration and visual interpretation of transcript isoforms, we developed IsoAtlas, an interactive database for examining both annotated and user-supplied transcript structures. Users can search known genes and transcripts or submit novel transcript models, visualize their structures, and assess their prevalence within indexed samples in IsoAtlas, expression, tissue distribution, disease context and sequencing-platform support across population-scale long-read RNA-sequencing data. A distinctive feature of IsoAtlas is its integrated human-mouse framework, which links corresponding transcript models across species and enables users to determine whether an isoform observed in a model organism is also supported in humans, and in which tissues or biological contexts. IsoAtlas also maps structurally equivalent transcripts across GENCODE, RefSeq and CHESS, providing simultaneous evidence for a transcript structure across different identifiers and reference annotations. The underlying evidence is derived from uniformly processed human and mouse long-read RNA-sequencing datasets indexed using Isopedia(14). Whereas Isopedia provides the underlying command-line framework for indexing and querying long-read transcript evidence, IsoAtlas extends this framework into an integrated database that combines interactive visualization, harmonized sample metadata, cross-reference transcript mapping, user-supplied transcript interrogation and human to mouse isoform relationships. Together, these capabilities provide a unified environment for visually exploring transcript diversity, interpreting novel isoforms and assessing the relevance of mouse transcript models to human biology. Unlike existing transcript databases that primarily catalogue transcript models, IsoAtlas contextualizes transcript isoforms using population-scale long-read sequencing evidence. IsoAtlas is freely available at https://www.isoatlas.org/.

## Results

### Overview of IsoAtlas

IsoAtlas is a scalable online resource that enables users to interpret arbitrary full-length isoform models against population-scale long-read RNA sequencing evidence, regardless of whether the transcripts are included in reference annotations as known transcripts or absent from them as novel transcripts (**Figure 1A**). IsoAtlas addresses this gap by establishing uniform population-scale evidence for transcripts together with standardized sample metadata. For known transcripts, IsoAtlas provides a cross-identifier mapping framework that connects equivalent transcript identifiers across commonly used reference annotations and establishes corresponding relationships between human and mouse transcripts. For novel transcripts, IsoAtlas accepts user-supplied files in GTF format. A gene query returns the unified transcript repertoire across GENCODE, RefSeq, and CHESS, together with transcript structures, sample prevalence, expression, tissue and disease context, and corresponding mouse transcript models when available.

**Figure 1.**
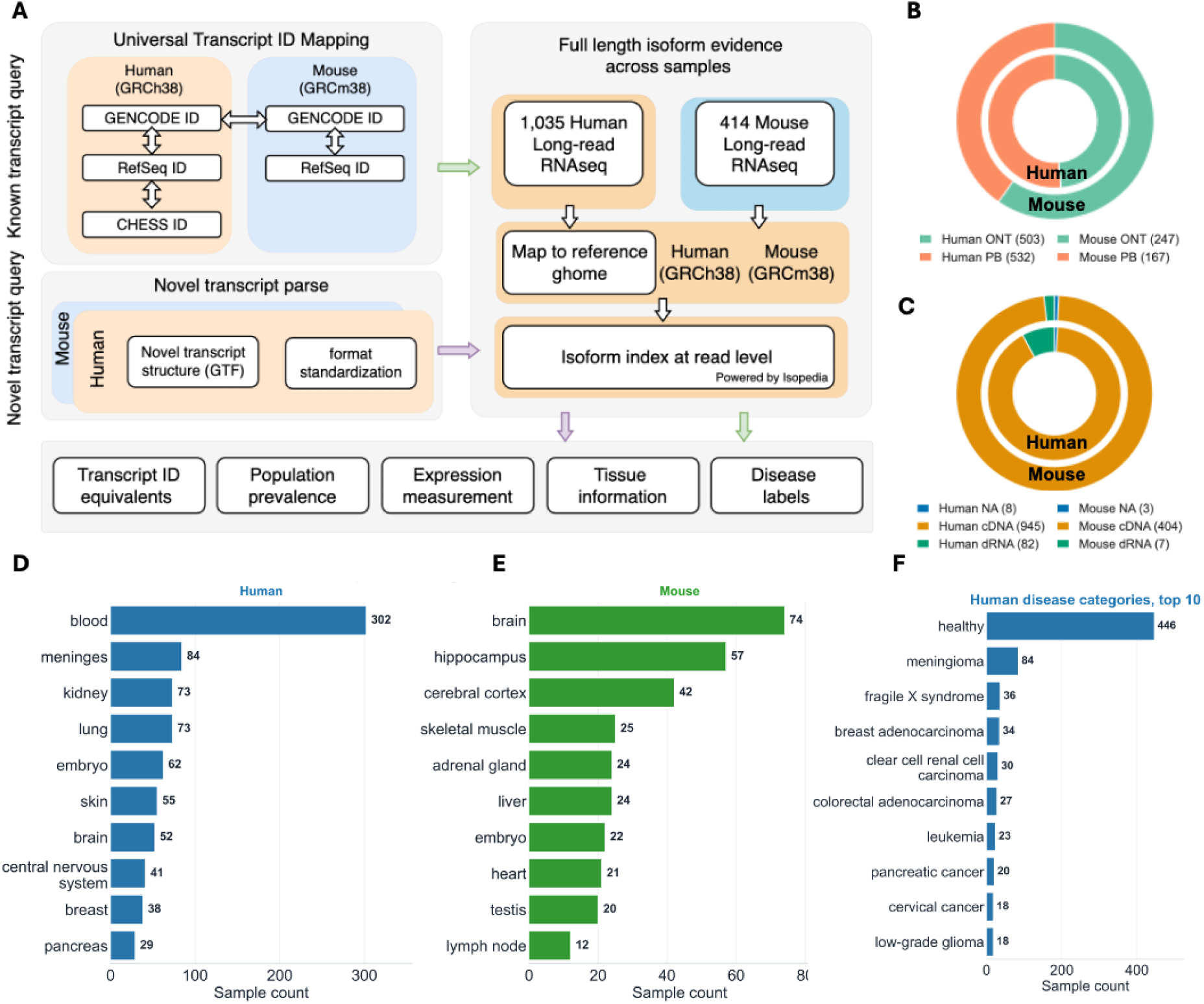
Overview of IsoAtlas. A, Schematic of the IsoAtlas architecture and workflow. Known transcripts can be queried through cross-annotation and cross-species identifier mapping, whereas novel transcripts can be submitted as GTF files. Long-read RNA-sequencing datasets are processed using a unified pipeline. IsoAtlas returns transcript-level information, including population prevalence, cross-reference transcript identifiers, expression measurements, tissue provenance, and disease labels. B–C, Sequencing platforms and chemistries represented in the human and mouse datasets, respectively. Inner circles for human samples, and outer circles for mouse samples. D-E, The ten most frequently represented tissues in IsoAtlas, with human tissues shown on the left and mouse tissues on the right. F, Distribution of disease labels across the human datasets.

The IsoAtlas backend currently comprises 1, 035 human and 414 mouse long-read RNA sequencing samples (**Supplementary Table S1** and **S2**). All human and mouse data were obtained from publicly available resources, including ENCODE(15) and the Sequence Read Archive (SRA)(16). These datasets were processed using a uniform workflow, beginning with read alignment, followed by the extraction and indexing of isoform signals using Isopedia, a command-line tool that enables rapid population-scale querying of transcript models across large sample collections (**Figure 1A**). IsoAtlas subsequently returns population-scale evidence, supporting reads in each sample, and the corresponding sample metadata.

Among the samples included in the current version of IsoAtlas, 750 were generated using Oxford Nanopore Technologies and 699 using PacBio sequencing (**Figure 1B**). Most datasets were generated using cDNA-based protocols, whereas direct RNA sequencing accounted for only 6% of the samples (**Figure 1C**). In total, IsoAtlas contains 9, 269, 810, 640 read-level records, of which 6, 838, 380, 289 were derived from human samples and 2, 431, 430, 351 from mouse samples. The metadata for each dataset were manually curated, standardized, and mapped to relevant ontology systems. The resulting resource covers 51 tissues, where blood (302 samples, 29.2%) represents the largest tissue category among human samples and brain (74 samples, 17.9%) is the most highly represented tissue among mouse samples (**Figure 1D** and **1E**, respectively). Furthermore, IsoAtlas includes 50 different disease labels that are currently available only for human samples, with healthy samples (446 samples, 43.1%) forming the largest category, followed by meningioma (84 samples, 8.1%) and fragile X syndrome (36 samples, 3.5%) (**Figure 1F**).

To establish an identifier-mapping framework spanning multiple annotation resources and species, we used IsoMatch(17) to merge transcript sets from GENCODE, RefSeq, and CHESS for human, and from GENCODE and RefSeq for mouse. Reference transcripts merged into the same transcript model were considered equivalent transcript models, thereby enabling the correspondence of transcript identifiers across different annotation resources. To further support cross-species studies, including those in which a biological mechanism is first characterized in a mouse system and subsequently translated to humans, we used the human-mouse transcript projections from ExTraMapper (18) to construct mappings involving 64, 233 human and 63, 951 mouse reference transcripts. When a transcript from one species is queried, the corresponding transcripts from the other species are displayed alongside it whenever a cross-species mapping is available.

### Cross reference query for known transcripts

Known transcripts catalogued in reference annotations form the basis of most transcriptomic analyses(2, 19). Commonly used resources such as GENCODE, RefSeq and CHESS provide complementary transcript collections that differ in coverage, curation strategy and intended use. As a result, the same transcript structure may be represented by different identifiers across annotations or may be included in one resource but absent from another, influencing how newly observed transcripts are classified(6, 7). IsoAtlas integrates these complementary annotations by comparing transcript structures and establishing correspondence between equivalent models across annotation systems. The current version of IsoAtlas integrates GENCODE v49, RefSeq 110, and CHESS 3.1, yielding 464, 936 non-redundant transcripts. Of these, 35, 956 (7.73%) are represented in all three reference annotations, 79, 854 (17.18%) are represented in two annotations, and 349, 126 (75.09%) are unique to a single reference annotation. IsoAtlas provides an easy to use interface that summarizes the queried gene or transcript (**Figure 2A**) and visualization for those known transcripts (**Figure 2B**). Based on this unified mapping framework, when users query any known transcript in IsoAtlas, the system simultaneously identifies its equivalent transcripts in other reference annotations. Because these transcripts represent identical or equivalent transcript structures, their population-level detection rates and sample-level expression measurements in IsoAtlas are linked and shared, thereby preventing supporting evidence from being fragmented because of differences in transcript nomenclature or annotation source.

**Figure 2.**
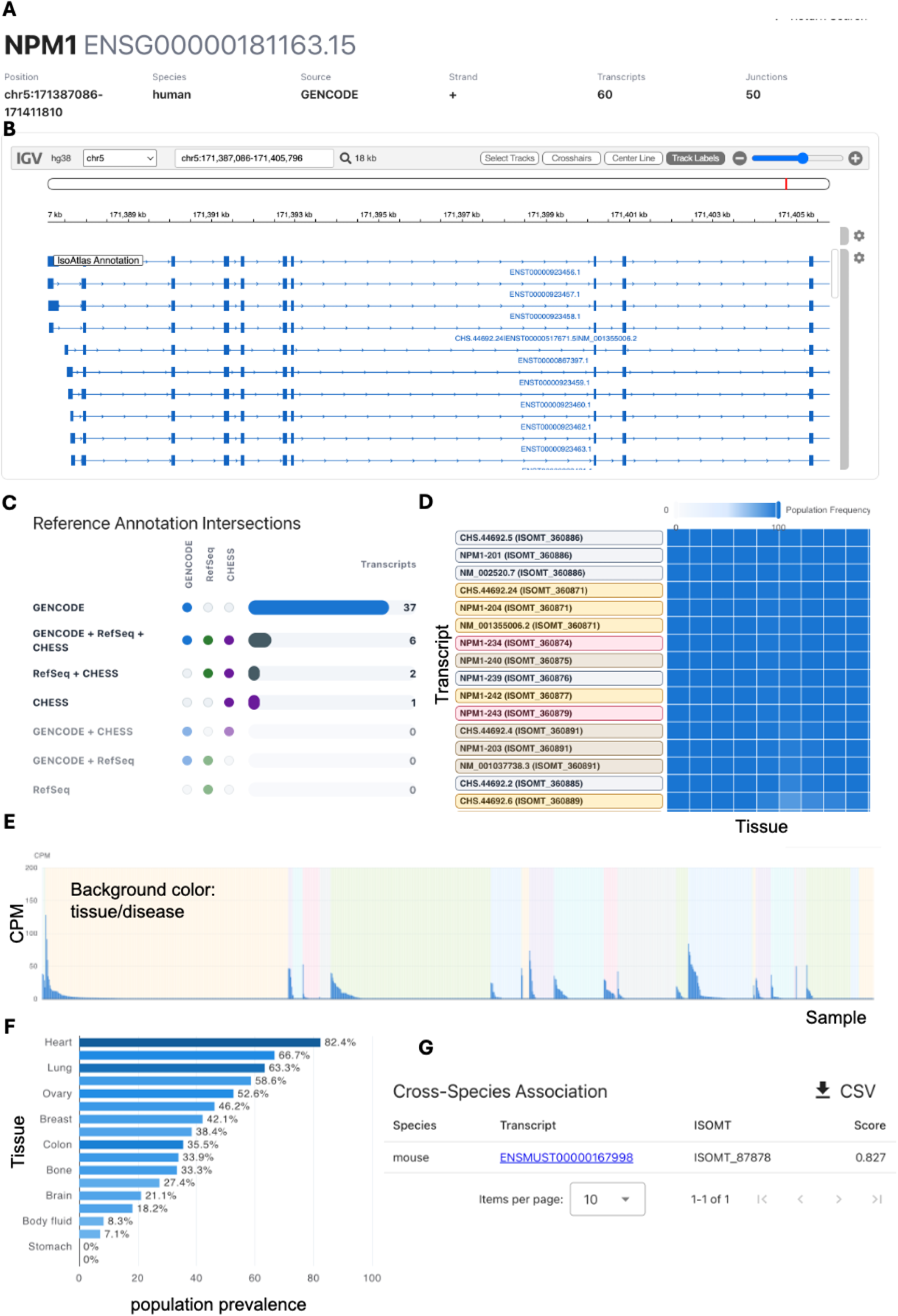
Web interface of IsoAtlas. A, IsoAtlas returned results summary B, Transcript structure browser supporting visualization of both known and novel transcripts. C, Cross-reference annotation mapping. When a known transcript is queried, structurally equivalent transcripts from other reference annotations are displayed simultaneously. D, Heatmap summarizing the population prevalence of all transcripts associated with a gene. Rows represent transcripts labeled with their original transcript identifiers, whereas columns represent tissues or disease categories. E, Transcript expression levels, measured as counts per million (CPM), across all samples. Samples are first grouped by tissue or disease category and then ordered by expression level in descending order. F, Detailed population prevalence information for an individual transcript. G, Cross-species transcript mapping. When a transcript has a corresponding transcript in another species, such as human or mouse, IsoAtlas reports the cross-species relationship on the transcript details page.

At the gene level, IsoAtlas displays all transcripts catalogued for a given gene across the different reference annotations and summarizes their overlap relationships in both tabular and UpSet plot formats (**Figure 2C**). The gene page also systematically summarizes the detection rates and expression levels of all known transcripts across samples and tissues. Expression is reported as counts per million reads mapped (CPM), facilitating comparisons of the population prevalence and tissue-expression characteristics of different transcripts from the same gene(**Figure 2D**). IsoAtlas provides a dedicated detail page for each individual transcript. Users may access this page either from the corresponding gene page or directly through the search function. The transcript detail page presents the transcript’s inclusion and mapping relationships across reference annotations, as well as its corresponding transcript relationships in other species. It further integrates information on population-level detection rate, expression level (**Figure 2E** and **2F**), tissue distribution, and disease labels, providing a unified entry point for the systematic interpretation of known transcripts and for comparisons across annotations and species.

### Cross-species querying facilitates the translation of mouse studies to human biology

Mouse models are central to functional genomics and studies of disease mechanisms(20). By linking human isoforms to corresponding mouse transcript models, IsoAtlas enables users to examine whether a human transcript structure is conserved and expressed in relevant mouse tissues. This information can support transcript prioritization and assess whether the corresponding isoform structure is represented in the mouse and thereby inform the suitability of the mouse model for isoform-specific studies..

To support this application, IsoAtlas integrates transcript-level relationships between human and mouse from the ExTraMapper database, enabling cross-species queries for known transcripts. As shown in **Figure 2G**, when a human transcript is queried, the corresponding gene page summarizes the number of transcripts within that gene that have associated mouse transcripts. In addition, the “Per-transcript summary” table lists the mouse transcript corresponding to each human transcript. Users can follow the associated link to open the corresponding mouse transcript detail page and further examine its transcript structure, population prevalence, expression level, and sample provenance.

### Novel transcript queries enable interpretation of transcripts outside reference annotations

Novel transcripts absent from all reference annotations generally lack stable identifiers, which precludes identifier-based queries. IsoAtlas circumvents this limitation by accepting a user-supplied GTF file for structure-based interrogation (**Figure 2A**). The submitted annotation is first validated and standardized; the corresponding Isopedia index is then selected according to the specified species, and each transcript structure is queried on the fly. As for known transcripts, IsoAtlas reports the prevalence of every queried structure across the 1, 449 long-read RNA-sequencing samples, together with the associated sample information.

Results for the entire submission are consolidated on a main result page, where transcripts are organized into tables and heatmaps that display sample prevalence alongside expression level, enabling direct comparison across the submitted set. Each transcript is further linked to a dedicated detail page reporting its prevalence across IsoAtlas samples, its sample-level expression, and the corresponding sample metadata, including tissue of origin and disease label. This structure-based query framework thus affords transcripts lacking reference annotation or standardized identifiers the same population-scale evidence and biological context that IsoAtlas provides for annotated transcripts.

### Case study of IsoAtlas for transcripts in gene TP53

To demonstrate the practical utility of IsoAtlas, we showcase it on TP53, well-established tumor suppressor gene frequently implicated in cancer (21, 22).

We first explored the reference transcripts cataloged in IsoAtlas based on its TP53 gene page. Among all 26 non-redundant known transcripts of TP53 from the reference annotations, five are included in all three annotations. Three are shared by RefSeq and CHESS, and the remaining 18 are annotated only in GENCODE. **Figure 3A** shows this information from the IsoAtlas visualisation. These results demonstrate that substantial differences in reference transcript content remain among commonly used annotations, even for a gene as extensively studied as TP53. By further integrating sample disease metadata, we found that reference transcripts exhibit distinct population provenance. This is shown on the isoAtlas panel as **Figure 3B**. For example, ENST00000269305.9, ENST00000620739.4 and other transcripts sharing the universal transcript ID ISOMT_177486 show the highest population frequency (71.88%) and are detected across multiple tissues, whereas transcripts such as CHS.21217.15 (CHESS) at 48.79% and NM_001126118.2 (RefSeq) at 36.62% display different population frequency levels.

**Figure 3.**
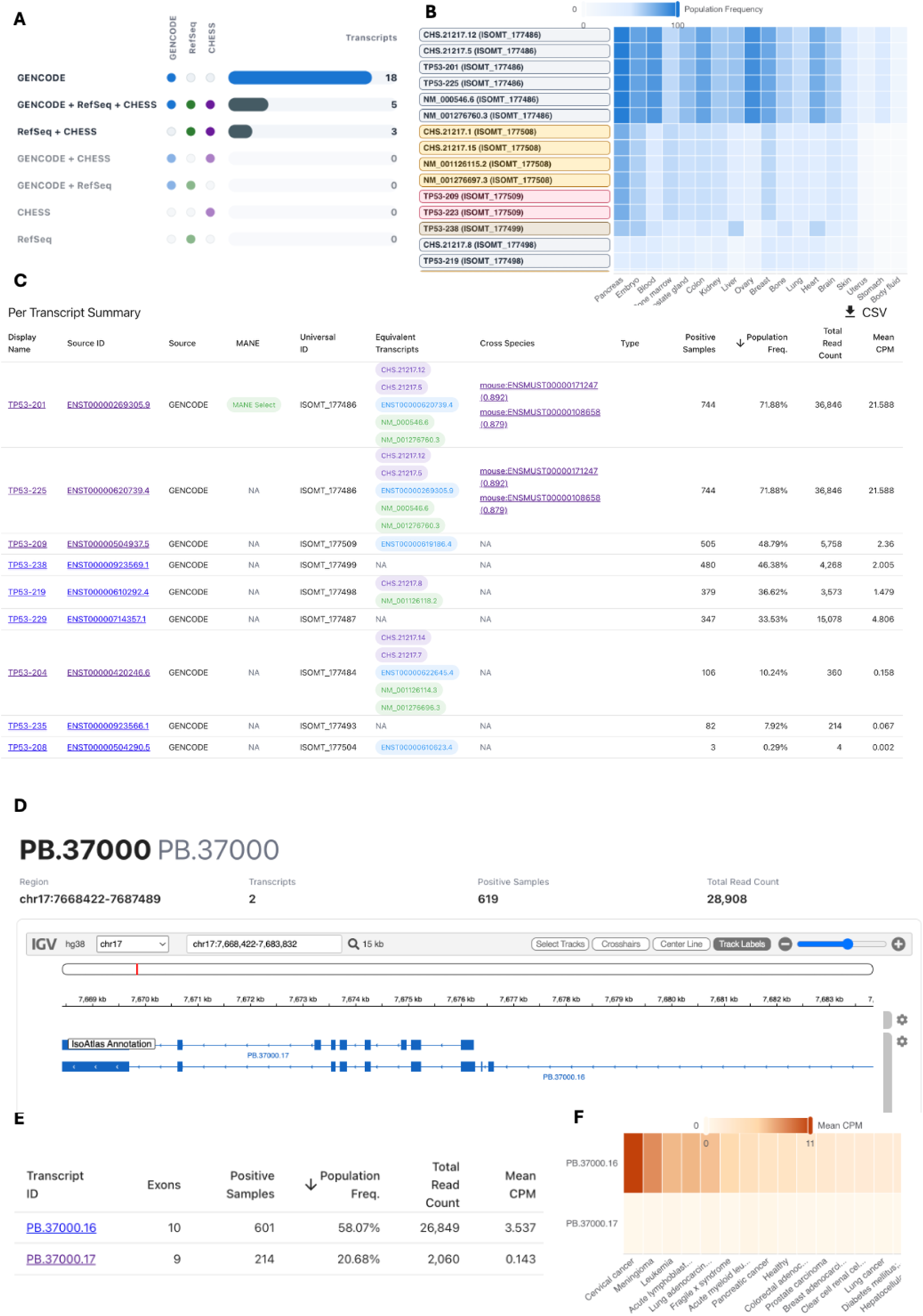
Population-scale characterization of known and novel TP53 transcripts using IsoAtlas. A, Intersection and distribution of known TP53 transcripts across the GENCODE, RefSeq, and CHESS reference annotations. B, Representative TP53 reference transcripts and their population frequencies across disease groups. C, Known TP53 transcripts summary from the HG002 sample in IsoAtlas. D, IGV visualization of novel transcripts. E, Population prevalence and mean expression levels (mean CPM) of two novel TP53 transcripts identified from the HG002 PacBio dataset. F, Comparison of the expression levels of the two novel TP53 transcripts across different disease groups.

To demonstrate IsoAtlas use case when analyzing a new sample, we collected TP53 transcripts identified from the HG002 cell line from long-read RNA PacBio sequencing dataset (see **Methods**). For HG002, we identified 27 TP53 transcripts. When comparing the 27 transcripts to the GENCODE v49 as reference annotation using IsoMatch, we identified 12 full splice match (FSM), 10 incomplete splice match (ISM), 1 novel in catalog (NIC) and 1 novel not in catalog (NNC) and 3 genic. Transcripts classified as NIC (PB37000.16) and NNC (PB37000.17) were considered novel transcripts. All FSM and ISM transcripts identified in HG002 were mapped to nine known transcripts as multiple transcripts from HG002 mapped into one reference transcript (**Supplementary Table S3**). The prevalence and expression levels of these nine known transcripts varied substantially, with population prevalence ranging from 0.29% to 71.88% and mean expression ranging from 0.0017 to 21.59 CPM.

Among these, transcript TP53-201(ENST00000269305.9) and TP53-202 (ENST00000620739.4) were linked to the mouse transcripts ENSMUST00000171247 and ENSMUST00000108658, respectively. These cross-species mappings suggest that functional studies of the corresponding mouse transcripts may provide insights into the biological roles of their human counterparts. Although TP53-201 and TP53-202 have identical transcript structures, they differ in their annotated coding sequences (CDSs). IsoAtlas therefore represents them as a single universal transcript structure and assigns them the same population prevalence and CPM values. This universal transcript structure showed the highest population prevalence (71.88% in humans and 63.77%–79.47% in mice) and expression levels (mean CPM: 21.59 in humans and 4.56–33.78 in mice) (**Figure 3C**).

We next focus on the two novel TP53 transcripts in the HG002 sample (**Figure 3D**). Using the custom transcript query interface in IsoAtlas, we found that both transcripts were recurrently detected across independent population samples. NIC transcript (PB37000.16) and NNC transcript (PB37000.17) showed population frequencies of 58.07% and 20.68%, respectively (**Figures 3E** and **3F**). In addition, PB37000.16 showed a mean expression of 3.537 CPM across the human IsoAtlas collection, compared with 0.143 CPM for PB37000.17. Thus, neither transcript appears to be specific to HG002. PB37000.16 was broadly detected across the IsoAtlas collection, whereas PB37000.17 was less prevalent but was nevertheless supported in 214 independent samples. These results place both transcripts within the broader spectrum of recurrent human TP53 isoforms rather than as isolated transcriptional products of the HG002 sample. Stratification by disease label showed that PB37000.16 was most frequently detected in cervical cancer datasets (**Figure 3F**). Although this pattern requires further evaluation, it highlights a specific candidate for follow-up. Thus, rather than leaving 27 TP53 transcript models as an undifferentiated catalog, IsoAtlas placed nearly all of them into annotation or population context and reduced the result to a single transcript warranting focused investigation.

## Conclusion and perspectives

IsoAtlas provides an interactive framework for exploring and interpreting transcript isoforms beyond the boundaries of any single reference annotation. By combining population-scale long-read evidence with transcript visualization, cross-annotation mapping and human-mouse transcript relationships, IsoAtlas enables users to determine whether known or user-supplied isoforms are recurrent, tissue- or disease-associated, supported across sequencing platforms and represented in a relevant model organism. This is particularly valuable for novel isoforms, for which absence from a reference annotation provides little information about prevalence or biological context. IsoAtlas therefore complements established transcript catalogs such as GENCODE, RefSeq or CHESS by adding an evidence layer that connects transcript structure to population prevalence, expression and sample provenance. IsoAtlas will be maintained at isoatlas.org and updated as additional public long-read RNA datasets and reference annotation releases become available. Each release will be versioned, and prior database versions and associated metadata tables will remain archived. Population prevalence adds a missing dimension to isoform interpretation. Conceptually, this parallels the contribution of population allele-frequency resources to variant analysis, although transcript prevalence is not equivalent to allele frequency. Rather than relying only on a binary known-versus-novel classification, IsoAtlas enables users to determine whether a transcript structure is recurrent, restricted to a particular tissue or disease context, or rarely observed across the current collection. These estimates should be interpreted as contextual evidence rather than as hard filtering thresholds, because transcript detection depends on expression, tissue representation, sequencing depth and library design. As additional datasets, tissues, cell types and species are incorporated, IsoAtlas will provide increasingly refined evidence for transcript recurrence, biological context and cross-species conservation.

Together, IsoAtlas provides a practical foundation for moving from transcript discovery toward population-aware and cross-species interpretation.

## Materials and methods

### IsoAtlas database organization

IsoAtlas is organized around a layered data model that connects transcript structures, reference transcript identifiers, sample metadata, transcript-sample evidence, population summaries, ontology mappings and cross-species transcript relationships.

The central unit in this organization is a non-redundant transcript structure. Reference transcript identifiers from different annotations are treated as mappings to this shared structural entity rather than as independent evidence units. This separates the biological structure being queried from the annotation-specific names used to refer to it, allowing equivalent identifiers to share one consistent population evidence record.

IsoAtlas therefore separates relatively static reference information from evidence-derived information. Static layers include transcript definitions, identifier mappings, sample metadata, ontology mappings and cross-species relationships. Evidence layers include transcript-sample support and aggregated population statistics. This separation keeps the database non-redundant while allowing different query types to join the relevant layers as needed.

For known genes and transcripts, IsoAtlas resolves the submitted identifier to the corresponding non-redundant transcript structure and retrieves the associated precomputed evidence, metadata and population summaries from SQLite3 databases. This supports rapid queries for reference-backed transcripts without repeatedly invoking the read-level query engine. See **Precomputation and query of known transcript evidence** section for details.

Novel transcript structures are handled through a separate dynamic path. Because user-supplied transcript structures cannot be exhaustively represented in the precomputed database, IsoAtlas queries the corresponding species-specific population index at runtime using Isopedia v1.6.8, then summarizes the returned evidence within the same transcript-sample and population framework used for known transcripts. See **Structure-based query of novel transcripts** section for details.

### Data collection and uniform processing

Publicly available human and mouse long-read RNA-sequencing datasets were collected from ENCODE and the Sequence Read Archive (SRA). ENCODE datasets were obtained from the ENCODE website, whereas SRA datasets were downloaded using the SRA Toolkit. Multiple sequencing runs originating from the same biological sample were consolidated at the sample level.

Reads were uniformly aligned using minimap2 (23) to GRCh38 for human and mm10 for mouse. Platform-specific parameters were `-ax splice:hq -uf` for PacBio and `-ax splice -uf -k14` for ONT. SAMtools (24) was used to sort the alignments and convert them to BAM format.

Isopedia v1.6.8 (14) was used as the read-level indexing and query engine. The `isopedia profile` command generated one profile for each dataset, and profiles from multiple runs of the same sample were consolidated using `isopedia-tools merge`. The resulting sample profiles were combined using `isopedia merge` and `isopedia index` to generate species-specific population indices.

### Sample metadata curation and standardization

Sample metadata were collected from the corresponding ENCODE and SRA records, including experiment and sample identifiers, study, species, sequencing platform, tissue, disease status and publication year. Tissue and disease annotations were manually standardized across studies, and synonymous labels were consolidated under consistent terms. Where applicable, standardized terms were mapped to ontology identifiers using the EMBL-EBI Ontology Lookup Service (OLS) (25).

These standardized metadata were linked to individual samples and used to summarize transcript prevalence and expression across tissues and disease groups.

### Construction of the unified IsoAtlas transcript catalogue

IsoAtlas integrates GENCODE, RefSeq and CHESS for human and GENCODE and RefSeq for mouse (see **Data Availability**). RefSeq GFF3 files were converted to GTF using gffread:

`gffread -T input.gff3 > output.gtf`

All annotations were standardized by retaining transcripts on the main reference chromosomes and converting chromosome identifiers to a consistent UCSC-style nomenclature.

Structurally equivalent transcripts across reference annotations were identified using IsoMatch v0.6.0:

`isomatch merge -D 0 -A 0 -U 0 -o merged ref1.gtf ref2.gtf ref3.gtf`

The resulting tracking information was parsed to link transcript identifiers representing the same exon–intron structure to a single non-redundant IsoAtlas transcript entity. Original GENCODE, RefSeq and CHESS identifiers were retained and associated with this shared structural representation, allowing population evidence to be calculated once per transcript structure rather than independently for annotation-specific identifiers.

### Cross-species transcript mapping

Human–mouse transcript relationships were obtained from ExTraMapper repository (https://github.com/ay-lab/ExTraMapper). The resulting transcript-level mappings were incorporated into IsoAtlas and maintained separately from within-species transcript equivalence. Human and mouse transcripts therefore retain independent species-specific entities and population statistics, while all retained one-to-one and one-to-many cross-species relationships are stored for comparative queries.

### Precomputation and query of known transcript evidence

Population-scale evidence was precomputed for every non-redundant transcript structure in the unified IsoAtlas catalogue. Each transcript was queried against the corresponding human or mouse population index, and sample-level supporting read counts and expression measurements were linked to standardized sample metadata. Derived population prevalence and tissue- and disease-associated summaries were stored together with the corresponding sample records in SQLite3.

Because structurally equivalent identifiers from different reference annotations are linked to the same IsoAtlas transcript entity, they share the same underlying population evidence. When a known transcript identifier is submitted, IsoAtlas first resolves it through the transcript mapping table and retrieves the corresponding precomputed evidence and metadata. Gene-level queries retrieve and summarize all unified transcript entities associated with the queried gene. Cross-reference identifiers and available human-mouse transcript mappings are subsequently joined to the returned results.

### Structure-based query of novel transcripts

Transcripts absent from the incorporated reference annotations generally lack stable identifiers. IsoAtlas therefore supports direct structure-based queries using user-supplied GTF files.

Submitted GTF files are parsed, validated and standardized to the chromosome naming and coordinate conventions used by IsoAtlas. The specified species determines whether the human or mouse population index is queried. Each transcript is then interrogated directly according to its structure without requiring a corresponding GENCODE, RefSeq or CHESS identifier.

Unlike known transcripts, whose evidence is precomputed, user-supplied transcripts are queried on the fly using Isopedia v1.6.8. Supporting-read evidence returned from the population index is linked to standardized IsoAtlas sample metadata to calculate population prevalence, expression and tissue- or disease-associated summaries using the same framework applied to known transcripts.

### Definition of prevalence and detectability metrics

When IsoAtlas compares the reads in a sample with a queried transcript, it returns the number of matching supporting reads. Supporting read counts are derived from two forms of transcript compatibility. Reads for which all splice junctions exactly match the queried transcript are counted as full-splice-match support. Reads whose junction chain represents a subset of the queried transcript are probabilistically assigned using an expectation-maximization algorithm. In IsoAtlas, a sample with a read count greater than zero is considered detectable. All detectable samples together define the prevalence of the transcript in the population, and the population frequency is calculated on this basis.

### Web implementation

The IsoAtlas website uses a decoupled front-end and back-end architecture. The front end is implemented as a single-page application using Vue.js (https://vuejs.org/) and Vuetify (https://vuetifyjs.com/), with ECharts (https://echarts.apache.org/) used for interactive data visualization. Tables in IsoAtlas support column-based sorting by clicking the corresponding column headers. Figures are interactive and allow users to inspect detailed information for individual data points. Both tables and figures can be downloaded using the corresponding download buttons.

Back-end data queries are served through the FastAPI framework. For known transcript queries, IsoAtlas precomputes Isopedia results and stores transcript population prevalence together with the associated sample information in SQLite3 databases. SQLite3 is also used to store sample metadata, mappings among transcript identifiers from different reference annotations, and cross-species transcript identifier mappings. Queries of novel transcript structures are performed on the fly using the Isopedia v1.6.8 binary.

### HG002 data processing

The HG002 transcript set was downloaded from the official PacBio FTP site and converted from GFF to GTF using GffRead. IsoMatch v0.6.0 was used to classify transcripts against GENCODE v49:

`isomatch classify -g ref.gtf -r ref.fa -o classified hg002.gtf`

TP53 transcripts were selected for downstream analysis. Known transcripts were resolved against the unified IsoAtlas transcript catalogue, whereas transcripts absent from the incorporated reference annotations were evaluated using the structure-based novel transcript query workflow. Population prevalence, sample support and expression summaries returned by IsoAtlas were used for the TP53 case-study analyses.

## Funding

This research was supported by NIH (1R01HG011774-01A1, 1U01HG011758-01 and 1UG3NS132105-01).

## Conflict of interest

FJS receives support from PacBio, ONT and Illumina.

## Data availability

IsoAtlas is available online for free at https://www.isoatlas.org/ and does not require user registration. Figures and tables can be downloaded by clicking the button on the figure. The datasets integrated in IsoAtlas are available at **Supplementary Table S1**(Human) and **S2**(Mouse). GENCODE reference annotations are downloaded from https://www.gencodegenes.org/human/release_49.html (human) and https://www.gencodegenes.org/mouse/release_M25.html (mouse). RefSeq reference annotations are downloaded from https://ftp.ncbi.nlm.nih.gov/genomes/all/annotation_releases/9606/110/GCF_000001405.40_GRCh38.p14/GCF_000001405.40_GRCh38.p14_genomic.gff.gz (human) and https://ftp.ncbi.nlm.nih.gov/genomes/all/GCF/000/001/635/GCF_000001635.20_GRCm38/GCF_000001635.20_GRCm38_genomic.gff.gz (mouse). CHESS annotation file is downloaded from https://github.com/chess-genome/chess/blob/master/chess3.1.3.GRCh38.primary.gtf.gz (Human).

HG002 transcript was downloaded from https://downloads.pacbcloud.com/public/dataset/Kinnex-full-length-RNA/DATA-Revio-HG002-1/5-Pigeon/isoseq_transcripts.sorted.filtered_lite.gff

## Notes

https://www.isoatlas.org/#/

